# Smooth muscle-specific MMP17 (MT4-MMP) defines the intestinal ECM niche

**DOI:** 10.1101/2020.06.18.147769

**Authors:** Mara Martín-Alonso, Håvard T. Lindholm, Sharif Iqbal, Pia Vornewald, Sigrid Hoel, Mirjam J. Damen, A.F.Maarten Altelaar, Pekka Katajisto, Alicia G. Arroyo, Menno J. Oudhoff

## Abstract

Smooth muscle is an essential component of the intestine, both to maintain its structure and produce peristaltic and segmentation movements. However, very little is known about other putative roles that smooth muscle may have. Here, we show that smooth muscle is the dominant supplier of BMP antagonists, which are niche factors that are essential for intestinal stem cell maintenance. Furthermore, muscle-derived factors can render epithelium reparative and fetal-like, which includes heightened YAP activity. Mechanistically, we find that the matrix metalloproteinase MMP17, which is exclusively expressed by smooth muscle, is required for intestinal epithelial repair after inflammation- or irradiation-induced injury. Furthermore, we provide evidence that MMP17 affects intestinal epithelial reprogramming indirectly by cleaving the matricellular protein PERIOSTIN, which itself is able to activate YAP. Together, we identify an important signaling axis that firmly establishes a role for smooth muscle as a modulator of intestinal epithelial regeneration and the intestinal stem cell niche.

## INTRODUCTION

The intestinal epithelium consists of a single layer of cells that is important for the uptake of nutrients as well as for providing a barrier to protect from pathogens. During homeostasis, intestinal epithelial cells are all derived from LGR5+ intestinal stem cells (ISCs) that reside at the bottom of crypts (Barker et al. 2007). Upon injury, however, or after depletion of LGR5+ cells, LGR5-cells can rapidly regain LGR5 expression and thus dedifferentiate to repopulate the crypt bottoms (Tian et al. 2011; Murata et al. 2020). In addition, within this dedifferentiation process, an epithelial reparative state exists, which is fetal-like, and depends on reprogramming by YAP (Yui et al. 2018; Gregorieff et al. 2015), and is further characterized by markers such as SCA-1 and HOPX (Wang et al. 2019; Nusse et al. 2018; Yui et al. 2018)

Adult intestinal epithelial (stem cell) maintenance relies on a variety of niche factors such as WNTs, R-spondins (RSPOs), Bone morphogenic proteins (BMPs), and prostaglandins, all of which are expressed by mesenchymal cell subtypes (McCarthy et al. 2020; Miyoshi et al. 2017; Greicius et al. 2018b; Shoshkes-Carmel et al. 2018a; Roulis et al. 2020; Degirmenci et al. 2018). These mesenchymal cells reside in the mucosa near the epithelium. However, intestinal epithelial homeostasis does not solely rely on soluble niche factors. The mechanical or extracellular matrix (ECM) niche is an additional defining factor, for example, by modulating the mechanosensory HIPPO/YAP pathway (Yui et al. 2018; Gjorevski et al. 2016). In addition, growth factors can interact with, or be embedded within the ECM to modulate their activity. Smooth muscle cells are one of the most prevalent non-epithelial cell types throughout the intestine, yet, their role in providing niche factors or affecting the ECM niche is largely unknown. Nevertheless, smooth-muscle specific deletion of tumor suppressor genes can result in defective epithelial growth (Katajisto et al. 2008). Furthermore, it was shown that cells that originated from the smooth-muscle can migrate into the mucosa to aid after injury (Chivukula et al. 2014). Nevertheless, it is still largely undefined what role adult smooth muscle has in ISC maintenance or during repair after injury.

Matrix metalloproteinases (MMPs) are fundamental ECM regulators, both by modifying ECM components directly and by cleaving growth factors to control their ability to bind the ECM and to the cell (Page-McCaw, Ewald, and Werb 2007a; Kessenbrock, Wang, and Werb 2015; Martín-Alonso et al. 2015). MMPs can play various roles in the injured intestine and many MMPs are upregulated in inflammatory bowel disease, likely by the increase in immune cell populations such as neutrophils with high proteolytic activity or in endothelial cells (O’Sullivan, Gilmer, and Medina 2015; Esteban et al. 2020). Thus, inhibition of MMP activity may be an attractive therapeutic target for treating inflammatory bowel disease (Jakubowska et al. 2016; Esteban et al. 2020). Although the role of certain MMPs such as MMP2, MMP7, MMP9, and MT1-MMP are relatively well-studied, the role of other MMPs in intestinal biology is still largely unknown. Here, we show that MMP17, a membrane-bound MMP which is specifically expressed in smooth muscle, is important to maintain optimal ISC stemness during homeostasis, and preserve the regenerative capacity of intestinal epithelium.

## RESULTS

### Intestinal smooth muscle is a rich source of BMP antagonists

Based on recent indications that fetal intestinal muscle is a provider of niche factors (Czerwinski et al. 2020), we first separated smooth muscle tissue and epithelial crypts from adult mouse colon and performed RNA-seq (Fig.1a, 1b, S1a). We found little evidence that intestinal smooth muscle expressed niche factors such as WNTs and RSPOs, or growth factors such as epidermal growth factor (EGF), however, we did find that smooth muscle expresses high levels of factors associated with BMP signaling including *Grem1, Grem2,* and *Chrdnl1* (Fig. 1b, S1a). Fluorescent *in situ* hybridization (FISH) confirmed high levels of these factors in a muscle-specific manner, in particular, we found enrichment of these factors in the muscularis mucosae that resides in close proximity to the bottom of epithelial crypts (Fig. 1c).

**Figure 1.**
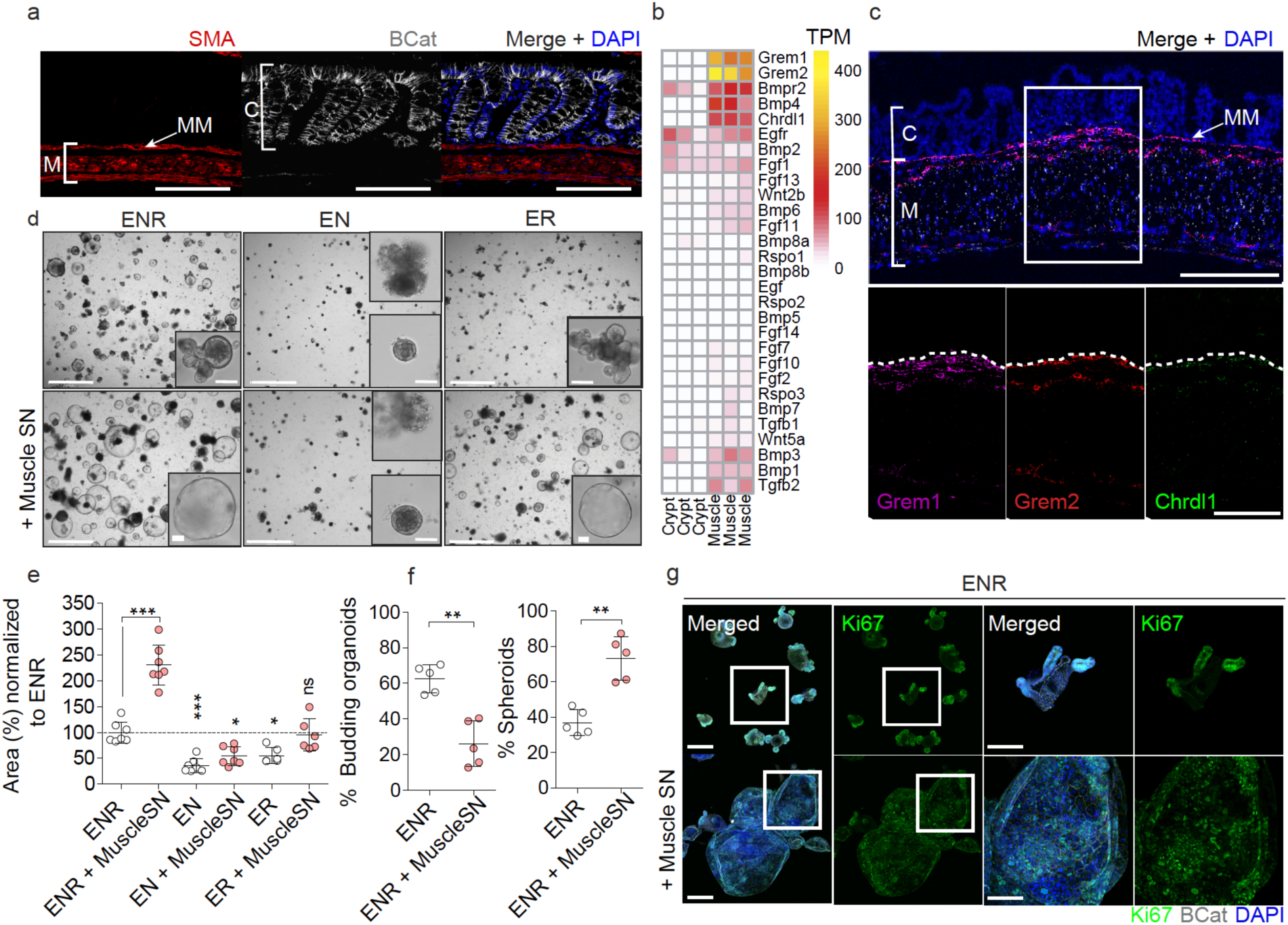
Intestinal Smooth muscle expresses BMP antagonists that control epithelial behavior. **a** Representative confocal maximum intensity projection of a transverse colon cut showing specific staining for muscle (M), SMA (Red), and crypts (C), βCatenin (βcat, Grey). Clear distinction among muscle layers can be observed, from muscularis mucosae (MM) close to the crypts to circular and longitudinal smooth muscle layers. Scale bar 100 µm. b. Heatmap depicting gene expression levels in TPM (Transcripts per million) in crypts vs muscle intestinal samples. n= 3 biological replicates. c. Representative confocal maximum intensity projections of fluorescence-coupled RNAscope showing BMP antagonists *Grem1* in magenta, *Grem2* in red, and *Chrdl1* in green, expressed in muscular cells. Scale bar 100 µm; 50 µm in magnified image. n= 2 mice. d. Representative bright field pictures showing organoids morphological changes when exposed to muscle-derived factors (Day4). Control media ENR (EGF, NOGGIN, RSPO1), EN (no RSPO1) or ER (no NOGGIN) were used to assess organoids reliance on external growth factors. Scale 1250 µm; 100 µm in magnified image. e. Graph shows organoids area in response to different media. Depicted average values normalized to ENR condition. n=5 to 7 independently performed experiments with 2-3 wells/experiment. f. Graphs represents the percentage of budding organoids or spheroids in response to ENR or ENR + muscle supernatant (Muscle SN). n=5 independently performed experiments with 2-3 wells/experiment. g. Representative maximun projection confocal images showing proliferative active cells (Ki67) in green and cell shape (βcat, grey) staining. Scale 200 µm, 100 µm in magnified image. n= 3 independent experiments. Numerical data are means ± SD. Data were analyzed by one-way ANOVA (e) or by Mann Whitney-test (f).

### Intestinal smooth muscle provides niche factors that render organoid growth independent of NOGGIN

Intestinal organoids are self-organizing *in vitro* epithelial structures that can be used to model *in vivo* processes (Sato et al. 2009; Serra et al. 2019). To test for a functional role of smooth muscle in providing niche factors, we exposed intestinal organoids to supernatant from muscle explants (muscle-SN) that completely lacked the mucosa (Fig. S1b). We found that organoids exposed to muscle-SN grew 2-3 times larger than in standard conditions and had a predominantly spheroid morphology (Figure 1d-g). Intestinal organoids rely on the supplementation of 3 factors epithelium itself does not express sufficiently; EGF, NOGGIN, and RSPO1 (ENR). Thus, we next tested whether muscle-SN could replace these factors. We found that muscle-SN was unable to replace RSPO1, as organoids failed to grow in RSPO1-deficient medium irrespective of muscle-SN supplementation (Figure 1d, 1e). In contrast, we found that muscle-SN was able to substitute for the BMP antagonist NOGGIN (Figure 1d, 1e). This is in support of our RNA-seq data in which we observed low *Rspo1/2/3* levels, but ample expression of BMP antagonists such as *Grem1* in smooth muscle tissue (Fig. 1b, 1c, S1a). ENR medium supports self-renewal and differentiation of organoids that acquire a typical structure including proliferative budding crypts and non-proliferative villus regions (Fig. 1f, 1g). In contrast, muscle-SN exposed organoids lacked crypts and consisted of mainly proliferative cells that were equally distributed along the spheroid (Fig. 1f, 1g).

### MMP17 is a smooth muscle-specific matrix metalloproteinase that is enriched in the muscularis mucosae

We find that intestinal smooth muscle expresses and secretes soluble niche factors that can mediate epithelial organoid growth (Fig. 1). Next, we wished to determine whether smooth muscle may also affect the ECM. The ECM acts on ISCs by providing a ‘mechanical’ niche as well as by being a reservoir of ECM-bound niche factors (Meran, Baulies, and Li 2017; Gattazzo, Urciuolo, and Bonaldo 2014; Kessenbrock, Wang, and Werb 2015). MMPs are known as important modulators of the ECM (Page-McCaw, Ewald, and Werb 2007b; Rodríguez, Morrison, and Overall 2010). By directly comparing epithelial crypt and smooth muscle we found several MMPs to be specifically expressed in smooth muscle tissue (Fig. 2a), including *Mmp17* (also known as MT4-MMP), a previously identified regulator of muscle growth factors and its surrounding ECM (Martín-Alonso et al. 2015). Therefore, we analyzed *Mmp17* expression in intestinal tissue by using the KO/KI mouse strain *Mmp17*^LacZ/LacZ^ (referred to as KO mice from hereon) (Rikimaru et al. 2007). We used LacZ staining (Fig. S2a) or a specific anti β-gal antibody to detect *Mmp17* promoter activity in the intestine from *Mmp17*^LacZ/+^ mice. As shown in Figure 2b, MMP17**^+^**cells are located only in the smooth muscle, and we identified that relatively more cells were MMP17**^+^** in the muscularis mucosae compared to the circular and longitudinal layers of the muscle (Fig. 2b, 2c). This is reminiscent of the expression of BMP antagonists (Fig. 1c), and indeed, using FISH, we find that *Mmp17* has an overlapping expression pattern with these BMP antagonists including enrichment in the muscularis mucosae (Fig 2d).

**Figure 2.**
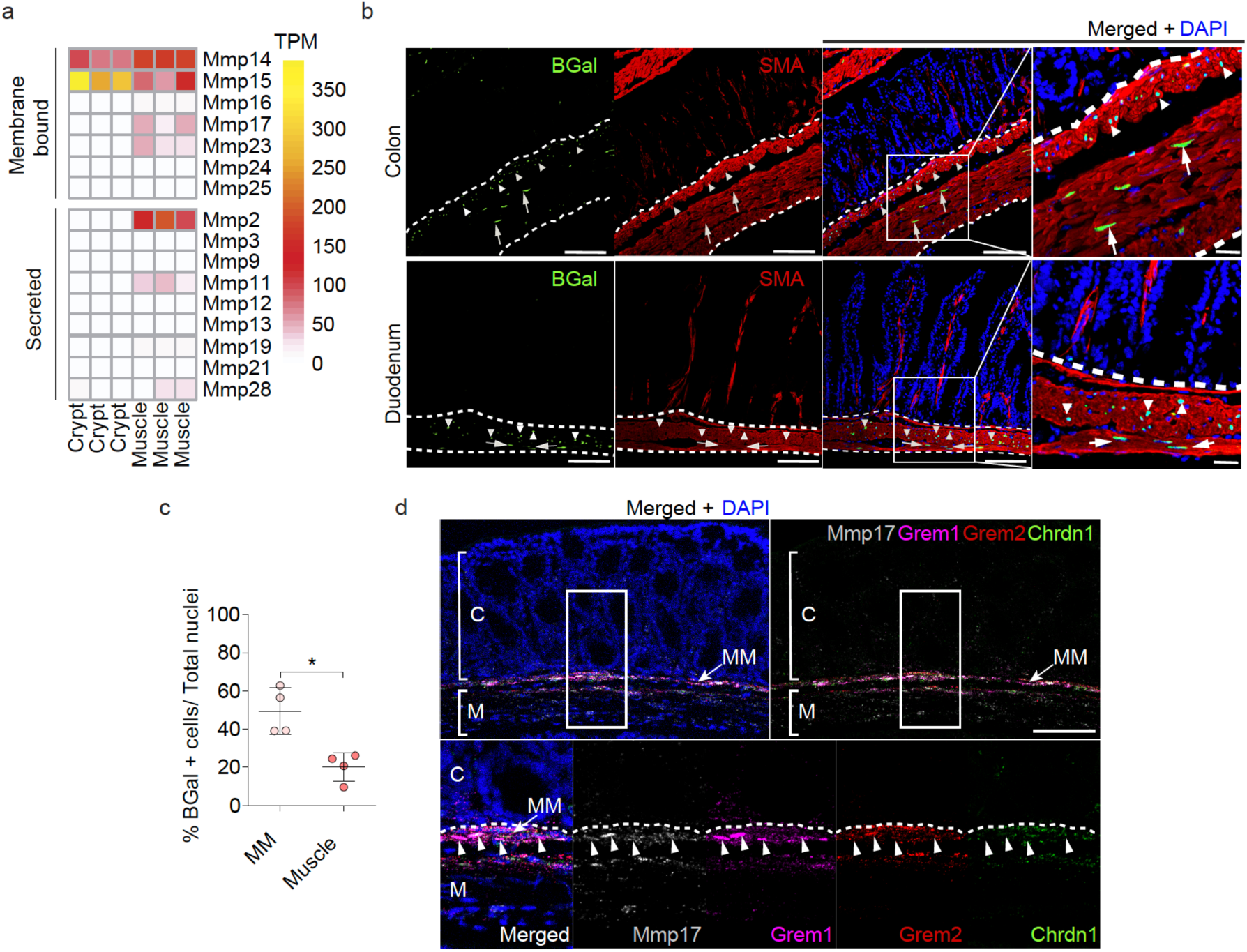
Muscle-specific matrix metalloproteinase Mmp17 is enriched in the muscularis mucosae together with BMP antagonists expressing cells. **a.** Heatmap shows TPM levels of Matrix Metalloproteinases (MMPs) in crypts vs muscle. n= 3 biological replicates. b. Representative confocal microscopy images showing active Mmp17 promoter (βGal staining, green) in positive SMA (red) muscle (white dashed line) stained in transverse intestinal sections of *Mmp17+/-* mice. Scale bar 100 µm; 20 µm in magnified image. n= 3 animals. c. Graph shows quantification of the % number of βGal positive cells in muscularis mucosae (MM) vs circular and longitudinal muscle (Muscle) in colon samples. n= 3 biological replicates. 4 images quantified. d. Representative confocal maximal projections of fluorescence-coupled RNAscope showing *Mmp17* (grey), and BMP antagonists (*Grem1*, magenta, *Grem2*, red and *Chrdl1*, green) co-expression in muscular cells (arrowheads). C, crypts, M, Muscle and MM, muscularis mucosae. Scale bar 100 µm; 50 µm in magnified image. n= 2 independent experiments with 1-2 samples/genotype. Numerical data in c are means ± SD and was analyzed by Mann-Whitney test.

### Muscle-specific MMP17 controls BMP signaling in crypts

Since MMP17 is enriched in the muscularis mucosa where its expression correlates with BMP antagonists, we asked about the possible impact of MMP17 loss in crypts. To unbiasedly gain mechanistic insight, we performed RNA-seq comparing WT and KO colonic smooth muscle and crypts. *Mmp17* is expressed in smooth muscle but not in the epithelium (Fig. 2), and we were surprised to find that only 42 genes were dysregulated in the KO muscle whereas 191 genes were dysregulated in KO crypts compared to their WT counterparts (Fig. 3a). In support, principal component analysis (PCA) was unable to distinguish WT from KO muscle, whereas KO crypts had a different distribution compared to WT crypts (Fig. 3b). Furthermore, *Mmp17* absence in intestinal smooth muscle did not result in any structural alteration at the smooth muscle level (Figure S3a). Upon closer examination, using the online gene enrichment tool Enrichr (E. Y. Chen et al. 2013; Kuleshov et al. 2016), we found SMAD4 as the top TF associated with upregulated genes in KO crypts (Fig. S3b). To test whether increased SMAD4 target genes in KO crypts was a direct result of SMAD4 protein levels, we performed immunostaining and western blot for SMAD4. Corroborating our unbiased transcriptome analysis, we found heightened nuclear localization of SMAD4 in the bottom of KO crypts compared to WT crypts, and increased levels of total SMAD4 in KO mucosa (Fig. 3d, 3e). The nuclear translocation or overall accumulation of SMAD4 is the result of cellular activation by TGFβ family members, including BMPs that specifically induce SMAD1/5/9 phosphorylation (Massagué 2012). Indeed, we found pSMAD1/5/9 to be particularly enriched in the bottom of KO crypts compared to WT crypts (Fig. 3f), which suggests altered BMP signalling in KO intestinal epithelium.

**Figure 3:**
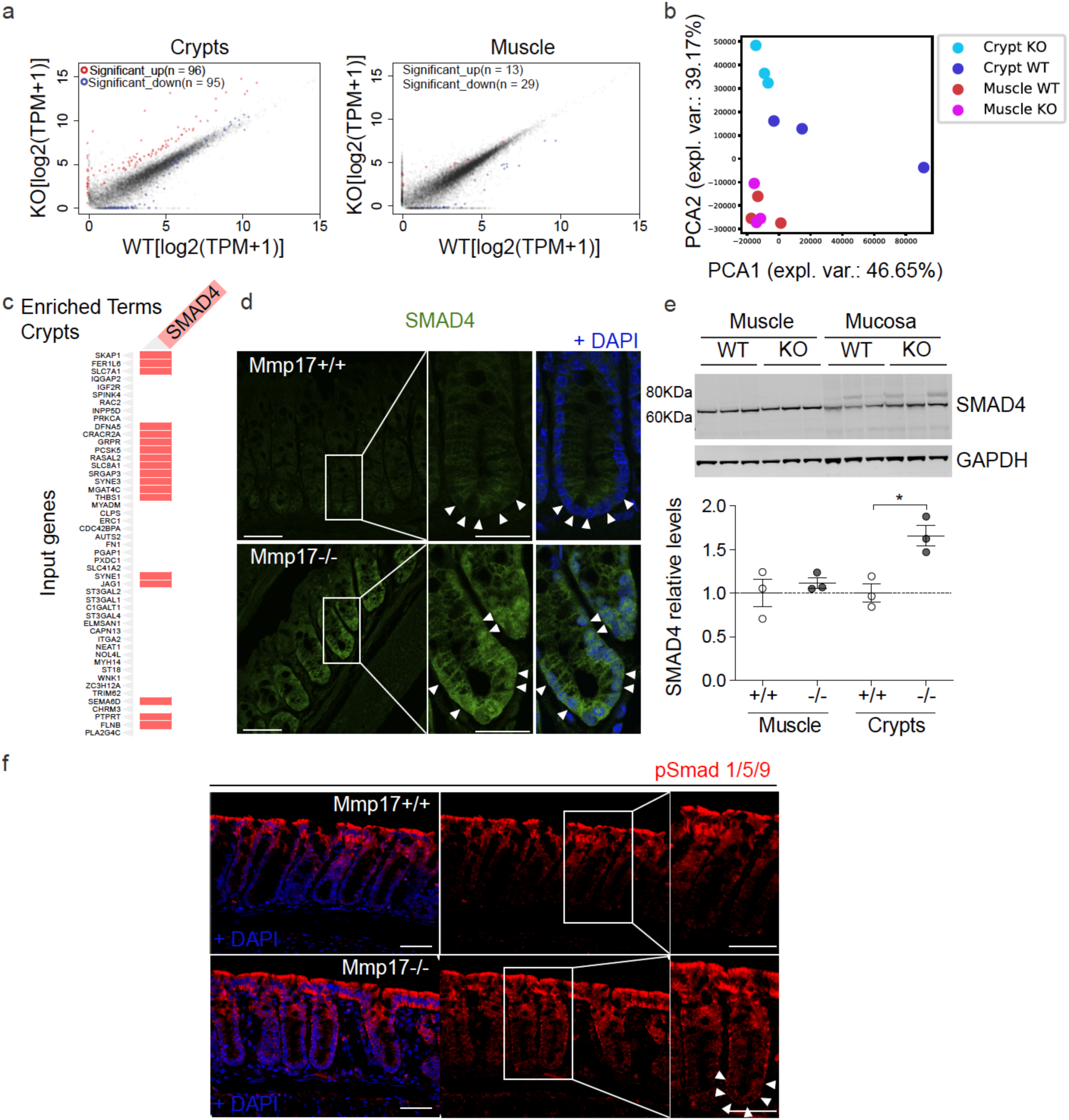
Muscle Mmp17 regulates crypts BMP signaling. **a** and **b.** RNA seq assay comparing WT and *Mmp17* KO crypts, we showed that loss of MMP17 in the muscle strongly affects epithelial gene expression to a greater extent than muscle gene expression. n= 3 biological replicates. c. Graph shows presence of genes related to SMAD4 signaling among the upregulated genes in WT vs KO crypts. d and e We observed higher SMAD4 signaling that was further confirmed by immunofluorescence in d (arrowheads indicate nuclear location in *Mmp17* KO) and western blot (e). n= 3 mice/genotype. Scale bar 50 µm, 25 µm in magnified views to the right. Representative confocal images showing pSmad 1/5/9 staining in intestinal crypts of WT vs KO mice. Arrowheads highlight crypt base pSmad 1/5/9 staining. Scale 50 µm. n= 2 biological replicates. Numerical data in e are means ± SD and were tested by one-way ANOVA followed by Tukey’s test.

### MMP17 regulates the ECM-ISC niche necessary for epithelial *de novo* crypt formation

SMAD signaling is essential to maintain ISCs, which is exemplified by the requirement of the BMP antagonist NOGGIN in intestinal organoid maintenance (Sato et al. 2009). To test if MMP17 affects ISCs, we quantified levels of the ISC markers *Lgr5* and *Olfm4* by ISH. We found that KO intestinal tissue had lower *Lgr5* and *Olfm4* levels compared to WT tissue in both the small and large intestines (Fig. 4a, 4b). In addition, decreased ISC marker gene expression was echoed by reduced organoid formation efficiency comparing KO with WT crypts (Figure 4c and Figure S4b). Importantly, organoid splitting, or culturing with excessive niche factors, resulted in equal organoid efficiency between WT and KO cultures (Fig. 4d). These data indicate that the reduced capacity of KO crypts to form organoids relied on the *in vivo* niche rather than a consequence of an intrinsic epithelial defect.

**Figure 4.**
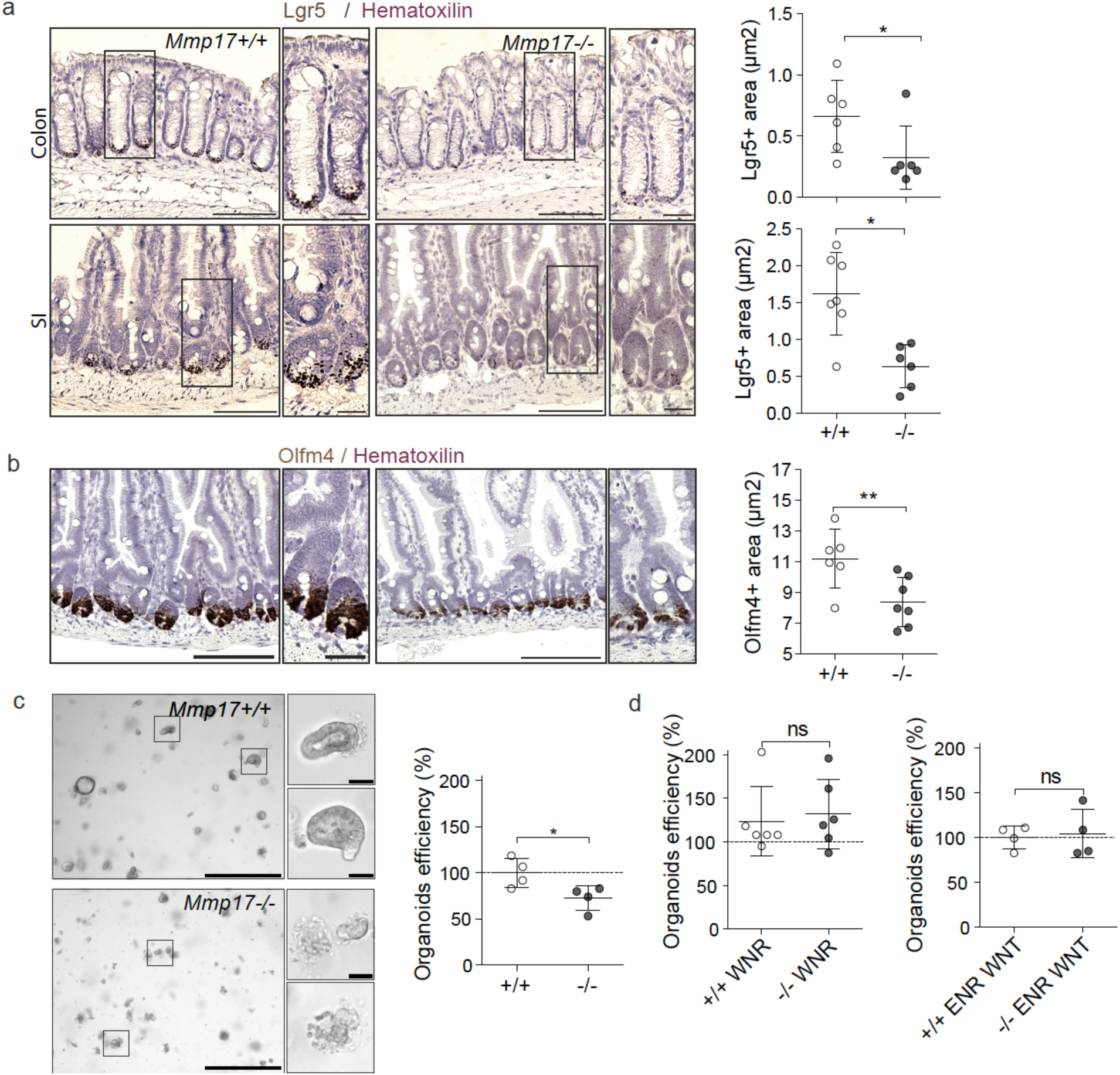
Muscular Mmp17 regulates crypt formation. **a-b.** Lgr5/Olfm4 RNAscope representative images of WT vs KO intestines and its quantification. Scale bar 100 µm; 25 µm in inset. n=6 mice per genotype analyzed in two different experiments. c. Representative bright field images of colon intestinal organoids from WT vs KO crypts 48h after crypt isolation (left). Scale 650 µm; 50 µm in cropped image. Graph shows relative organoids number as percentage normalized to WT. Each dot represents a well. One representative experiment is showed, out of 5 performed with similar results. d. Graphs represent the percentage of organoids derived from colon crypts after 72h in response to the enriched medium WNR (left) and this percentage after splitting (right). Numerical data are means ± SD and were tested by Mann-Whitney test.

### MMP17 is required for intestinal repair after inflammation or radiation-induced injury

*De novo* crypt formation is important during epithelial repair which is needed upon injury, for example after damage inflicted by inflammation or due to irradiation. To test if smooth muscle, and in particular MMP17, could play a role in intestinal injury responses, we used Dextran Sulfate Sodium (DSS) to induce experimental colitis and compare WT to KO littermates (Fig. 5a). At day 5, we found that both WT and KO mice had indistinguishable features of disease indicating that damage was induced equally (Fig. S5a-e). After 5 days, DSS was replaced by regular drinking water to allow for intestinal repair, which is initiated rapidly and requires reprogramming of the intestinal epithelium (Yui et al. 2018; Miyoshi et al. 2017). Two days after DSS, KO mice had shorter colons and sustained hemorrhage, including an increased presence of blood in stool compared to WT mice (Fig. 5b, 5c). Furthermore, other disease features were also exacerbated in KO mice compared to WT mice. We found that KO mice had a higher injury score than WT mice, as was determined using a genotype-blind injury classification based on H&E images (Fig. 5d, 5e). H&E analysis further revealed a larger area of ulcerated mucosa and lower presence of epithelial crypts in KO mice compared to WT mice (Fig. 5e). In addition, we found a significant reduction in proliferative epithelium in KO mice (Fig. 5f), suggesting that epithelial reprogramming towards a reparative state relies on MMP17.

**Figure 5.**
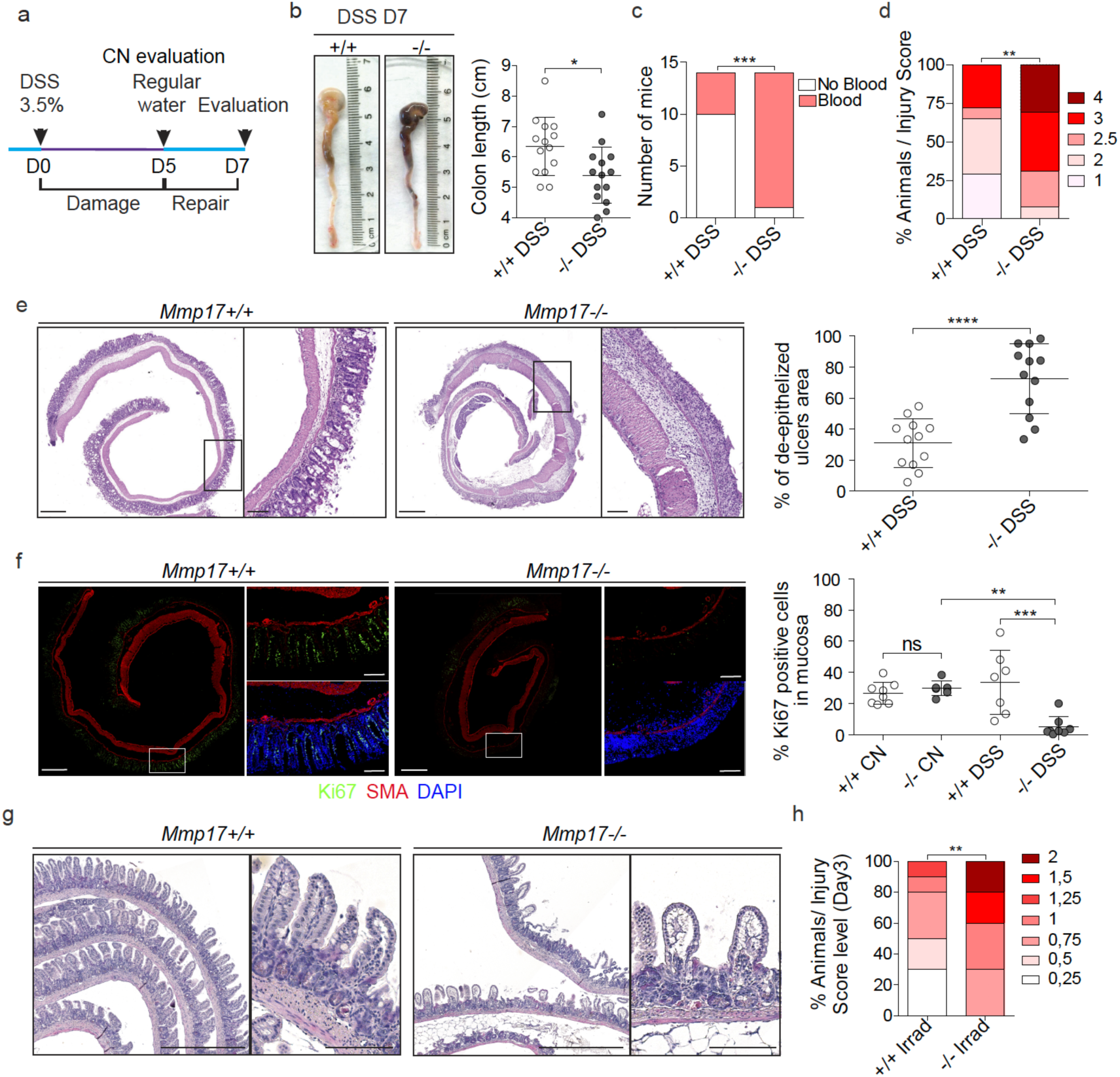
Muscle-specific Mmp17 is required for intestinal repair after injury. **a.** Timeline of DSS treatment. Mice were exposed to 3.5% DSS for 5 days followed by 2 days of regular water to allow epithelial restoration. Evaluation of control (CN) mice was performed at day 5 (D5). b. Representative images of mice colon in DSS-treated animals at D7. Graph shows colon length, and presence of blood in stool and/or colonic cavity (right) at end point (D7) in c. n= 14 animals per genotype, obtained from 5 independent experiments. d. Blind scoring of DSS-derived injury in colon, based on histological features. n= 14 mice per genotype from 5 independent experiments. e. H&E images showing distal colon in swiss roll. Scale 500 µm; 100 µm in magnified views to the right. Graph represents the damaged mucosa as % of area with ulcers relative to the total length of the distal colon. n= 12 mice per genotype. f. Representative maximal projection of confocal images of DSS-treated distal colon stained for the proliferative marker Ki67 (Green), SMA (Red) and DAPI (Blue). Scale 500 µm; 100 µm in magnified views. Graph shows % of proliferative cells (Ki67 positive) in mucosa, normalized to total mucosal cell number. n= 7-8 mice per genotype. g. Representative images of H&E sections of small intestinal swiss rolls 3 days after 10Gy irradiation. Scale 500 µm; 100 µm in magnified views. n= 7-8 mice per genotype analyzed from 2 independent experiments. h. Blind scored injury level in irradiated mice. n=7-8. Numerical data in b, e and f are represented as means ± SD. Data in b and e were tested by t-test. Data in c was analyzed by Fisher’s exact test (One-tailed) and in d and h were analyzed by Mann-Whitney test for nonparametric populations. Data in f was analyzed by one way ANOVA followed by Tukey’s Multiple Comparison test.

Next, we performed an alternative non-inflammatory injury model based on whole-body irradiation. Indeed, a single dose of ionizing radiation (10Gy) induces equal apoptosis in WT and KO intestines (Figure S5f). As custom in this model, WT animals regained crypt-villus structures 3 days after irradiation, however, small intestines of KO mice displayed disorganized epithelia with thicker and blunted villi (Figure 5g, S5g for quantification). In addition, a genotype-blinded evaluation of damage features in H&E images showed increased signs of damage in small intestines of KO mice compared to WT mice (Figure 5h). Together, these data indicate that muscle-specific MMP17 is required for appropriate intestinal epithelial reparative responses upon injury.

### *Mmp17* loss results in long-term reparative epithelial defects and increased tumor risk

We wondered whether intestines of KO mice would eventually heal, and thus we evaluated weight gain for a prolonged time after DSS administration (Fig. 6a, 6b). WT mice rapidly returned to their original weight; however, KO mice never fully regained their starting weight (Fig. 6b for females, Fig. S6a for males). In support, while WT mice largely returned to homeostatic conditions at end point, KO mice retained shorter colons, experienced sustained blood in stool, still had areas of unhealed ulcers in the epithelial surface and had higher injury score compared to WT mice (Fig. 6c-h). In addition, we detected the presence of crypt distortions named reactive atypia, and these morphological changes were predominantly found in KO mice (Fig. 6f, 6g, S6b). These morphological changes resemble those found in intestinal neoplastic lesions, so we next decided to evaluate the impact of *Mmp17* loss in the initiation and progression of intestinal tumors using the *Apc*^Min^ mouse model. *Apc*^Min^ mice develop tumors primarily in the small intestinal epithelium (McCart, Vickaryous, and Silver 2008). We found that loss of *Mmp17* predispose mice to the formation of a higher number of tumors both quantified macroscopically (Fig. 6i), and microscopically using H&E stained sections (Figure 6j). We did not observe differences in tumor area (Fig. S6c) suggesting that MMP17 mediates tumor initiation but not tumor progression. In support, WT and KO tumors were indistinguishable in terms of β-CATENIN and OLFM4 distribution (Fig. S6d). Of note, *Mmp17* expression was restricted to muscle cells also in tumor areas (Fig. S6e). In sum, our data indicate that MMP17 is required for short and long term intestinal epithelial repair and its loss predisposes to intestinal neoplastic alterations.

**Figure 6.**
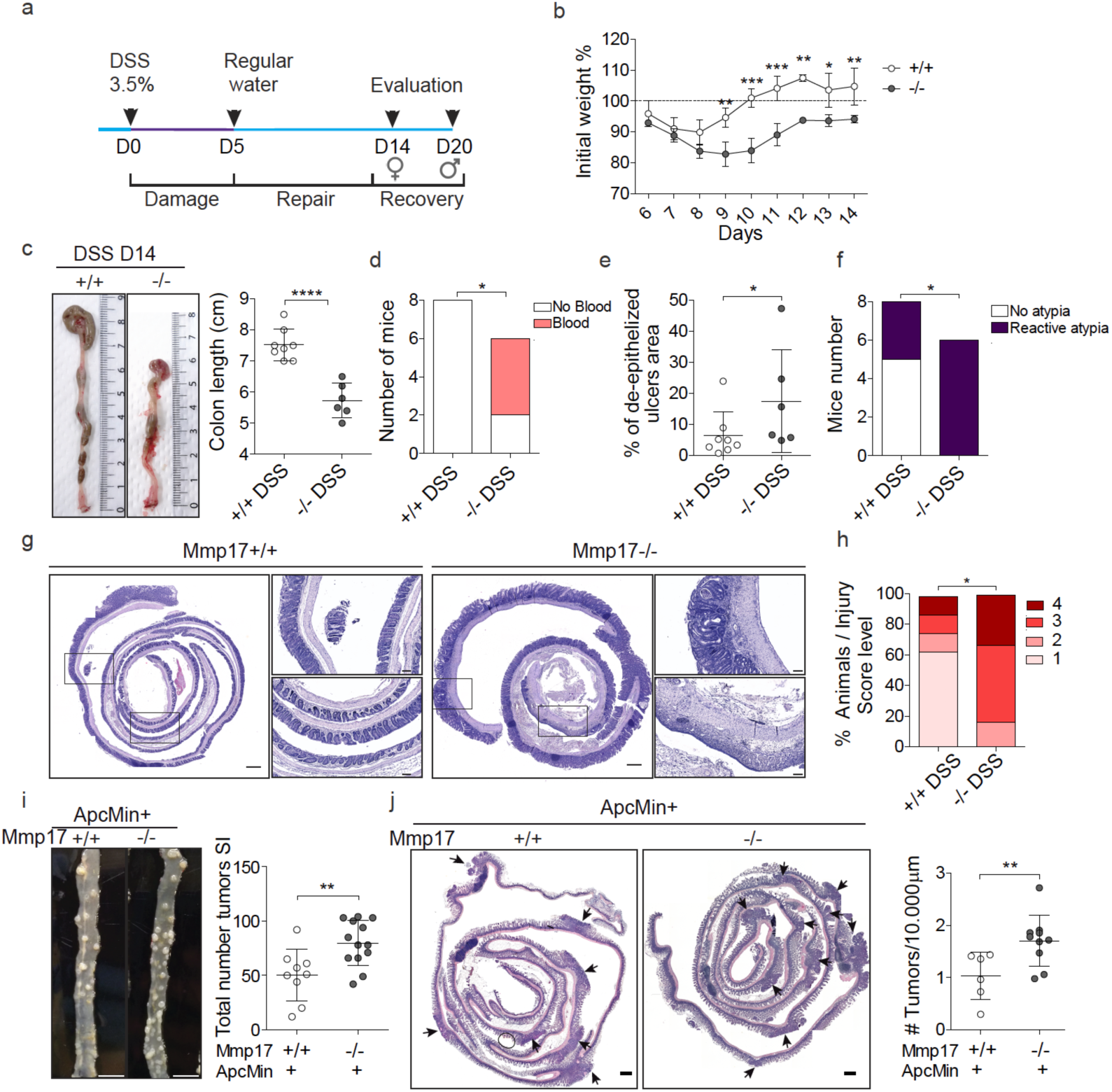
MMP17 absence hinders long-term repair in mouse intestinal epithelia and leads to increased tumorigenesis. **a.** Timeline of DSS-long treatment. Mice were exposed to 3.5% DSS for 5 days followed by 14 (females) or 19 days (males) days of regular water to allow epithelial restoration. b. Weight loss and recovery relative to % to initial weigh. n= 3 to 4 mice per genotype and gender. Represented one experiment with females. c. Representative images showing colon length at end point and its quantification. n= 6-8 mice. d. Graph shows presence of blood in stool/colon lumen at end point. n= 6-8 mice per genotype. e. Graph represents the percentage of unhealed areas (ulcers) in WT and KO colonic mucosa at long term. f. Graph shows incidence of reactive atypia (or reactive epithelial changes after DSS). g. Representative H&E pictures of colon swiss rolls showing healed crypts in WT vs ulcered areas and reactive atypia in KO. n= 6-8 mice per genotype. Scale 500 µm. h. Injury score representation of damaged colon evaluation. n= 6-8 mice per genotype. i. Representative pictures showing tumor incidence in a portion of the small intestine of *ApcMin+ Mmp17* WT and KO mice (Jejunum). Scale bar 1 cm. Graph shows total number of tumors counted in fresh tissue (small intestine complete length). n= 9 to 13 mice per genotype. j. H&E representative images of small intestine swiss rolls in transverse cut. Arrows highlight visible tumors. Scale bar 500 µm. Graph shows tumor quantification normalized to tissue length. n= 6 to 10 mice per genotype. Numerical data in b was analyzed by two way ANOVA followed by Bonferroni posttest. t test (normal distribution) was applied in c. Fisher’s exact test, one tailed was used in d and f. Numerical data in e and h were analyzed by Mann-Whitney t-test. Data in i, j were analyzed by unpaired t-test.

### Muscle-SN induces a reparative epithelial state that includes activation of YAP

Since we observed that smooth muscle MMP17 is essential for intestinal repair we wondered whether muscle indeed can promote epithelial reprogramming towards a reparative state. We found that muscle-SN can replace NOGGIN in the culture media, and muscle-SN treated organoids grow as large spheroids (Fig. 1). Organoid spheroid growth can either be characterized as reparative that is associated with fetal-like gene programs, such as organoids derived from SCA-1+ cells (Nusse et al. 2018), or it can be the result of increased WNT signalling such as upon treatment with the GSK3 inhibitor CHIR (Yin et al. 2014). These two different organoid spheroid states are on opposite ends when it concerns ISCs; a reparative state is characterized by a loss of LGR5+ ISCs whereas ISCs expand upon CHIR treatment (Yui et al. 2018; Nusse et al. 2018). To determine what type of spheroids are induced by muscle-SN, we performed RNA-seq comparing normal organoids to muscle-SN treated organoids (Fig. 7a). Using gene set enrichment analysis, we found that genes associated with LGR5+ cells, as well as genes upregulated in organoids treated with CHIR, were downregulated in muscle-SN treated organoids (Fig. 7b). In contrast, gene sets associated with repair and fetal programs were significantly enriched in muscle-SN treated organoids (Fig. 7b). Intestinal epithelial repair programs are coupled with YAP signalling and, indeed, a YAP signature gene set was also enriched in muscle-SN treated organoids (Fig. 7b). In support, we found that YAP was localized nuclear throughout muscle-SN induced spheroids, whereas it was cytoplasmic in the centre of budding organoids (Fig. 7c). Together, we conclude that muscle-SN induces spheroid growth that is associated with reparative/fetal-like reprogramming and may be mediated by YAP.

**Figure 7.**
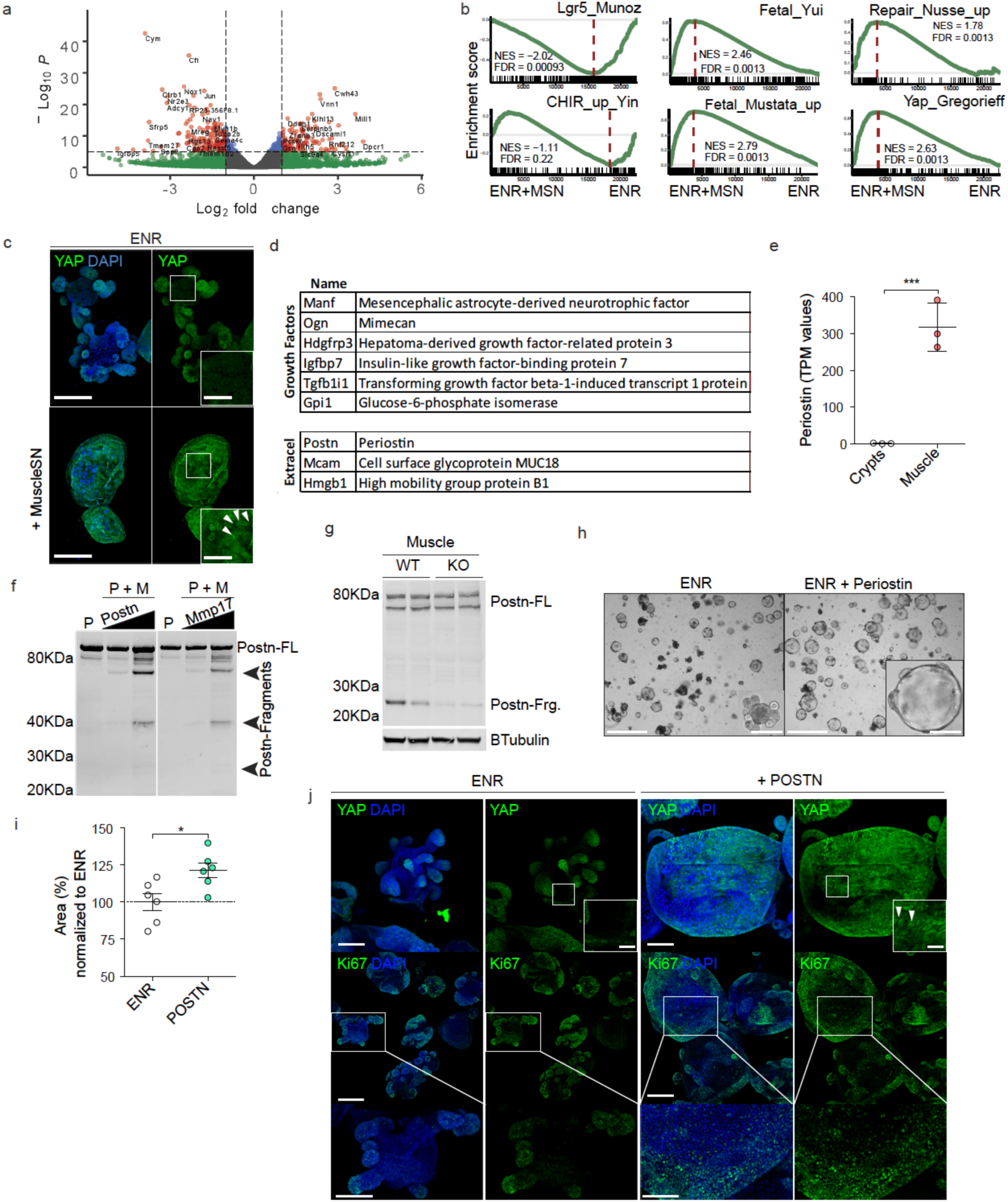
Identification of muscle-derived factor PERIOSTIN as an in vivo and in vitro substrate for MMP17. **a.** Volcano plot showing significantly up and downregulated genes in organoids exposed to muscle-SN vs ENR. b. GSEA of indicated gene sets comparing ENR with MuscleSN treated organoids. NES (normalized enrichment score) and FDR (false discovery rate) are depicted. c. Representative maximal projection confocal images showing cytoplasmic vs nuclear YAP staining (green). Scale 100 µm;25 µm in inset. Arrowheads highlight nuclear YAP. n= 2 independent experiments. d. List of growth factors and extracellular proteins found in muscle-SN. n= 3 biological replicates. e. TPM values for *Postn* comparing crypts with smooth muscle tissue. n=3 biological replicates. f. In vitro digestion experiment with human recombinant proteins showing POSTN fragments when in contact with MMP17 catalytic domain. P, POSTN, M, MMP17, FL, full length. n= 2 experiments with increasing concentrations of POSTN or MMP17. g. WB of mouse intestinal muscle showing decreased POSTN fragments in the absence of MMP17. n= 2 biological replicates. h. Representative brightfield images showing organoids morphology in the presence of POSTN and area quantification (Day 3) (i). Scale 1250 and 200 µm in inset. n= 3 independently performed experiments with 2-3 wells/experiment. j. Representative confocal maximum intensity projection images showing YAP and Ki67 (green) staining in POSTN-treated organoids. Scale 100 µm; 25 µm in inset (YAP pictures) and 200 µm and 100 µm in Ki67 pictures. Arrows highlight nuclear YAP. Numerical data are means ± SD. Data in e represents p-adjusted value from RNAseq analysis and data in i was analyzed by Mann-Whitney test.

### PERIOSTIN is a muscle-derived factor cleaved by MMP17 that activates YAP and induces organoid growth

MMP17 is expressed in intestinal smooth muscle and is required to allow for epithelial repair *in vivo* (Figs. 5, 6). Furthermore, muscle-SN is able to reprogram epithelium towards a reparative state *in vitro* (Fig. 7a-c). We therefore hypothesize that MMP17 cleaves a muscle-derived factor that facilitates epithelial reprogramming that is needed for repair. To identify said factor, we examined muscle-derived proteins by performing mass-spectrometry on muscle-SN, and we detected 550 proteins (Suppl. Table 1). We next curated this list to only display proteins known to be growth/niche factors and/or extracellular proteins (Fig. 7d). Among these was PERIOSTIN (POSTN), which is a matricellular protein that is highly expressed in smooth muscle (Fig. 7e). We previously identified a different matricellular protein, OSTEOPONTIN (OPN), as an MMP17 substrate (Martín-Alonso et al. 2015). Of note, POSTN can serve as a ligand for integrins to activate YAP signalling (Ma et al. 2020). Further, POSTN has been proposed to capture BMP members in the ECM (Maruhashi et al. 2010). Both of these features are relevant for biological processes we identified to be affected by MMP17. Co-incubation of POSTN with MMP17 led to several POSTN fragments indicating MMP17 is able to cleave POSTN (Fig. 7f). In addition, we examined POSTN in MMP17-WT and KO smooth muscle and we found a decrease in POSTN fragments in KO muscle samples, suggesting that cleavage of POSTN also occurs *in vivo* and it is impaired in KO intestines (Fig. 7g). Finally, we found that POSTN itself induces organoid growth that is associated with induction of Ki67 and nuclear YAP (Fig. 7h-j).

## DISCUSSION

The role of smooth muscle cells, other than in peristalsis, has been largely undefined. However, in a recent preprint, it was found that in early human gut development ACTA2+ muscularis mucosa cells are the major source of WNT, RSPO, and GREM niche factors (Czerwinski et al. 2020). In contrast, in adult mice, there have been various mucosa-resident mesenchymal cell populations described that express *Rspo* and *Wnt* genes (Greicius et al. 2018a; Shoshkes-Carmel et al. 2018b; Degirmenci et al. 2018). We here find that smooth muscle, and in particular, the muscularis mucosa, is the primary source of BMP antagonists such as *Grem1/2* (Fig. 1). The importance of GREMLIN1 as an ICS niche factor was recently determined by an experiment in which *Grem1*-expressing cells were depleted using the diphtheria-toxin system, which also led to the rapid loss of LGR5+ ISCs (McCarthy et al. 2020). This experiment would thus have led to the death of practically all muscularis-mucosa smooth muscle cells. Together, our work here as well as recent work by others highlight the importance of smooth muscle for providing niche factors important for ISC homeostasis.

We find an important role for MMP17 *in vivo*. We show that KO mice have a reduction of ISC-associated genes in homeostasis (Fig. 4), which may be caused by increased levels of SMAD4 at the bottom of KO crypts (Fig. 3). Indeed, BMP signalling can impair ISC-signature genes by direct repression *via* SMAD1/4 recruitment of HDAC1 (Qi et al. 2017). We further identify that MMP17 is required for intestinal epithelial repair after injury, which we link to the ability of MMP17 to cleave POSTN. We hypothesize that MMP17 cleavage of POSTN is necessary upon injury to reprogram epithelial cells in a YAP-dependent manner. Earlier this year, Ma *et al.* described a role for POSTN in activating YAP/TAZ through an integrin-FAK-Src pathway using cancer cell lines (Ma et al. 2020). We here extend these findings and show that POSTN can also affect primary non-tumor cells (Fig. 7). In addition to this direct role affecting the epithelium, others have shown that POSTN can alter the ECM by binding to a variety of ECM associated proteins including BMP1, FIBRONECTIN, and TENASCIN-C (Kudo and Kii 2018). We would speculate that through this link with the ECM, cleaved POSTN may broadly affect intestinal homeostasis *in vivo,* which would be impaired in MMP17-deficient mice such as we observe.

In addition, others have also shown that POSTN is cleaved *in vivo*, suggesting this has a physiological role, and *Postn*-deficient mice have general issues with repair in various tissues (Shimazaki et al. 2008; Nishiyama et al. 2011). Thus, although the importance of cleavage of POSTN remains not fully comprehended (Kudo 2011), it is clear that POSTN plays an important role in reparative processes and the identification of MMP17 as an enzyme that can cleave POSTN can be used in future studies.

MMP17, unlike other MMP members, exert minimal activity against classical ECM components (Kolkenbrock et al. 1999). So far, only a few bona fide substrates have been described in other tissue environments, such as ADAMTS-4 (Clements et al. 2011), alphaM integrin (Clemente et al. 2018) and two matricellular proteins; OPN (Martín-Alonso et al. 2015) and POSTN described in this work. We can not discard the possibility that these or other unidentified MMP17 substrates play a role in ISCs niche regulation or the intestinal response to injury, particularly OPN which expression is increased in patients with inflammatory bowel disease (Neuman 2012). Furthermore, in a tumor environment, both ADAMTS4 and OPN are overexpressed in colon cancers (M. Zhao et al. 2015; H. Zhao et al. 2018; J. Chen et al. 2018). Finally, also non-catalytic activities for MMP17 have been described related to tumors (Paye et al. 2014).

To summarize, we discover a previously unappreciated role for intestinal smooth muscle tissue. We find that smooth muscle are likely the major contributor of BMP antagonists, which are essential niche factors for the maintenance of ISCs. In addition, we describe an important role for smooth-muscle restricted expression of MMP17, which *in vivo* is required for epithelial repair. Finally, we provide evidence that MMP17 acts *via* cleavage of the matricellular protein POSTN, which in itself can induce repair-like features in intestinal epithelium.

## ACKNOWLEDGEMENTS

We would like to thank Dr. Motoharu Seiki for kindly sharing the *Mmp17*^KO/KI^ mouse line and Dr. Kaisa Lehti for first inspiring this study. We would like to thank to Anne Beate L. Marthinsen for performing the irradiation, and the Department of Radiology and Nuclear Medicine (St. Olav’s Hospital) for allowing the use of their instruments. We also thank Arne Wibe and Elin Rønne for their evaluation of colon crypt reactive atypia. We thank the imaging (CMIC) and animal care (CoMed) core facilities (NTNU), as well as the histology and animal facilities at CNIC for assisting in this work. The WT KO crypt-muscle RNA-seq was done by the Genomics Core Facility at NTNU, which receives funding from the Faculty of Medicine and Health Sciences and Central Norway Regional Health Authority. This work was further financially supported by the Norwegian Research Council (Centre of Excellence grant 223255/F50, and ‘Young Research Talent’ 274760 to MJO) and the Norwegian Cancer Society (182767 to MJO). MMA is the recipient of a Marie Skłodowska-Curie IF (DLV-794391).

## DATA DEPOSITION

All raw sequencing data is available through ArrayExpress: WT and KO smooth muscle and crypt RNA seq: E-MTAB-9180; ENR vs MuscleSN treated organoids RNA seq: E-MTAB9181.

## SUPPLEMENTARY FILE

**Figure S1.**
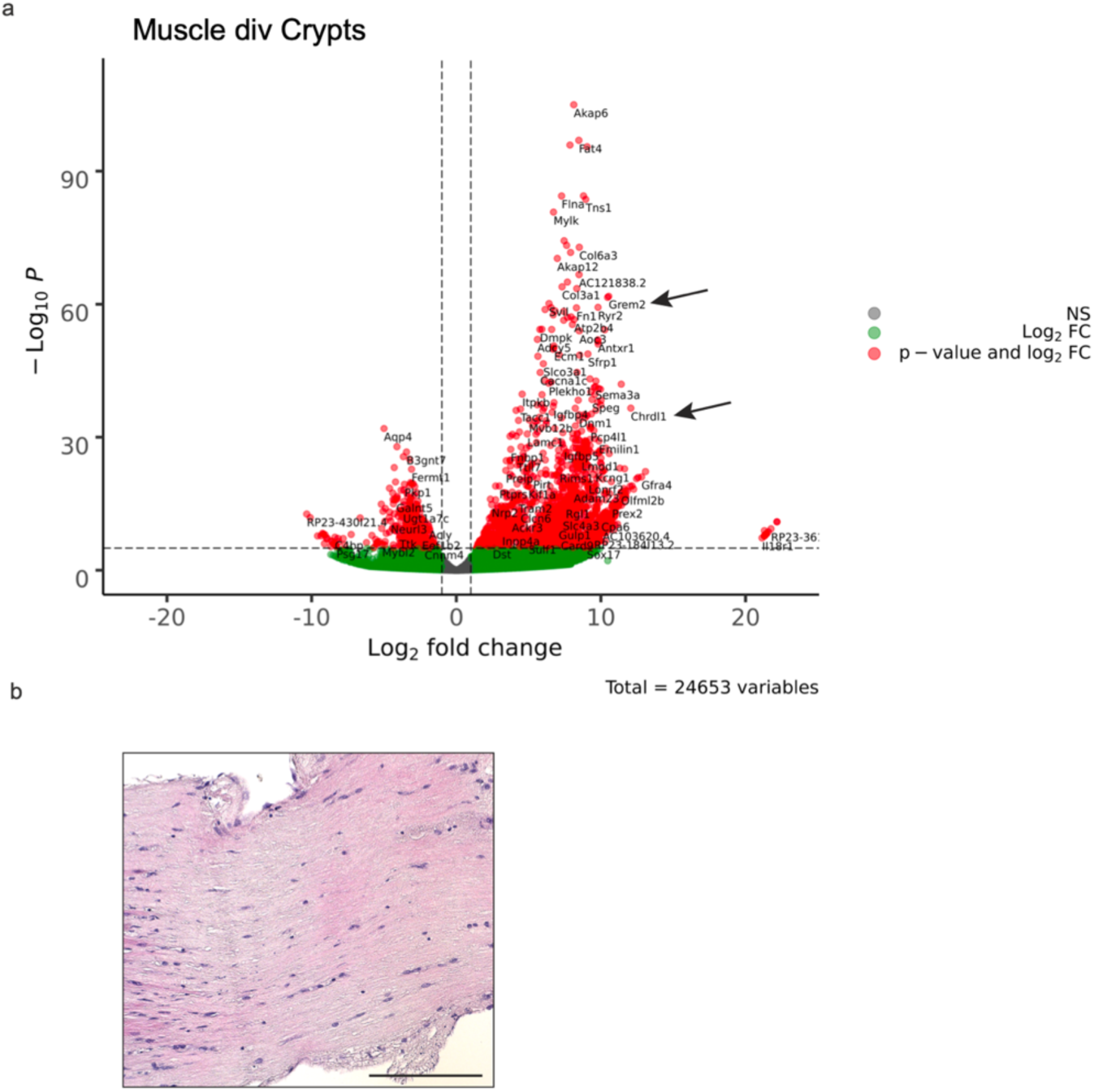
RNAseq of intestinal crypts vs muscle reveals the unique expression of growth factors by muscle. **a** Volcano plot showing differential expression of genes between muscle and crypts as Gremlins and Chordin like1 (arrows), BMP signaling antagonists. n= 3 biological replicates b. Image of H&E stained colon muscle strip used to obtain muscle-SN (24h after isolation). Scale 150 µm. n= 3.

**Figure S2.**
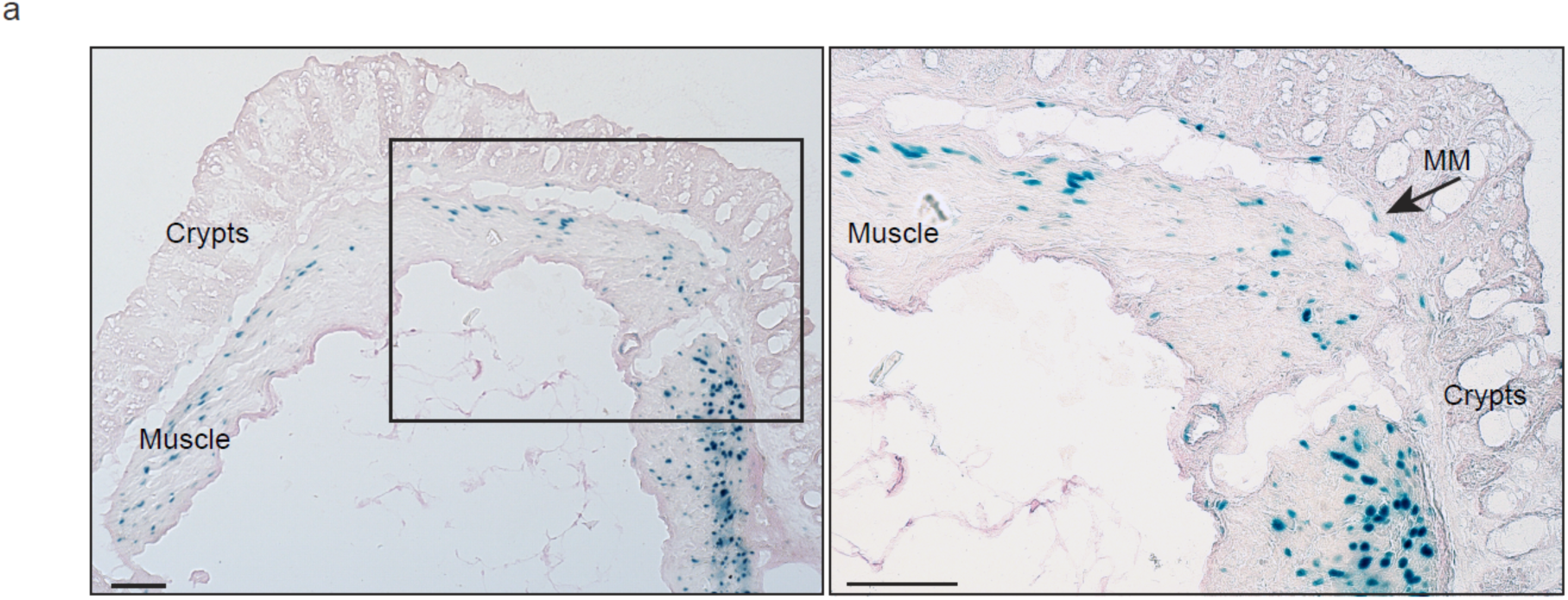
Mmp17 promoter is active in muscle cells from muscularis mucosa and circular and longitudinal muscle. **a.** Representative image of a transverse colon cut stained for β-Galactosidase activity (blue). Scale bar 100 µm.

**Figure S3.**
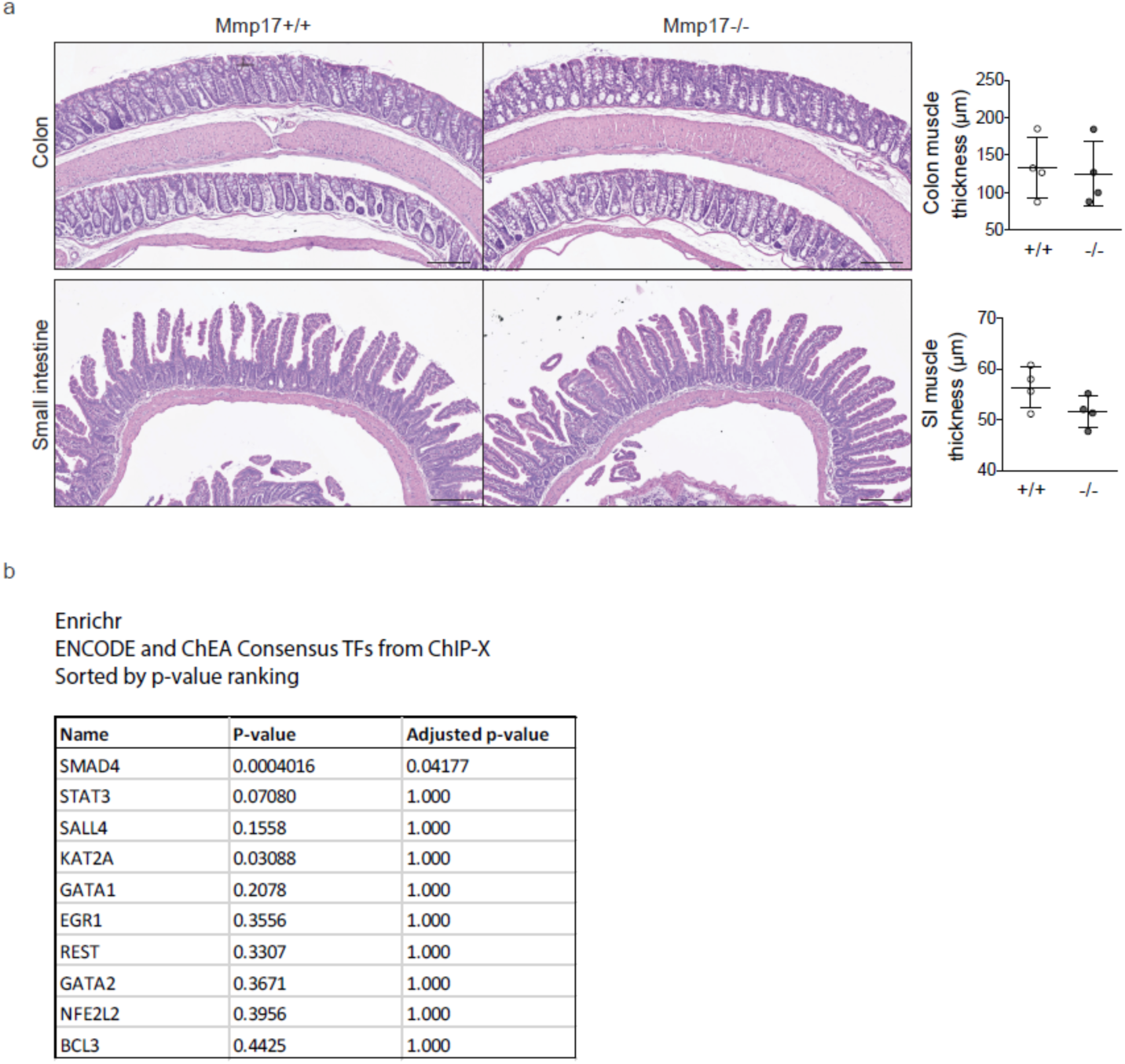
No structural differences in KO muscle and ENRICHR results on transcription factors differences between WT and KO crypts. **a.** Representative H&E images of transverse colon and small intestine (SI) tissues. Scale 150 µm. n= 4 mice per genotype. Graphs represent average values for muscle thickness along the swiss roll. b. List of transcription factors altered when comparing WT and KO epithelium (Enrichr). Data in a was analyzed using Mann-Whitney test.

**Figure S4.**
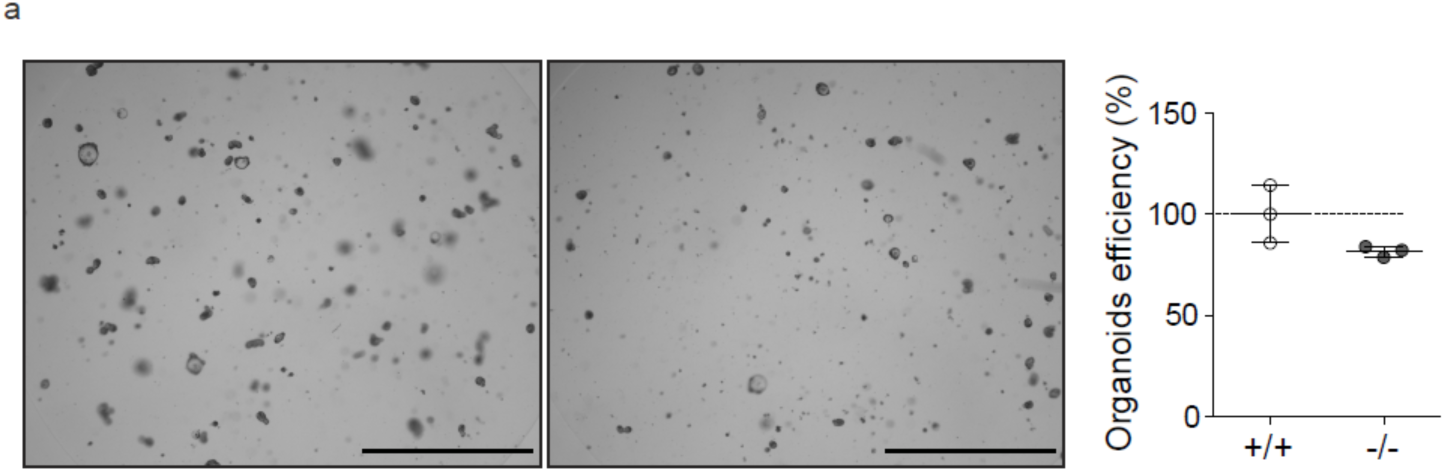
Lower levels in stem cell marker Olfm4 and lower organoids efficiency formation in Mmp17-/- small intestine. **a.** Bright field representative images of small intestine organoids 72h after crypt isolation. Graph represents organoids efficiency after 72h of WT or KO crypt culture. n= 3 wells analyzed. Scale 650 µm. Data in a are means ± SD and were analyzed using Mann-Whitney test.

**Figure S5.**
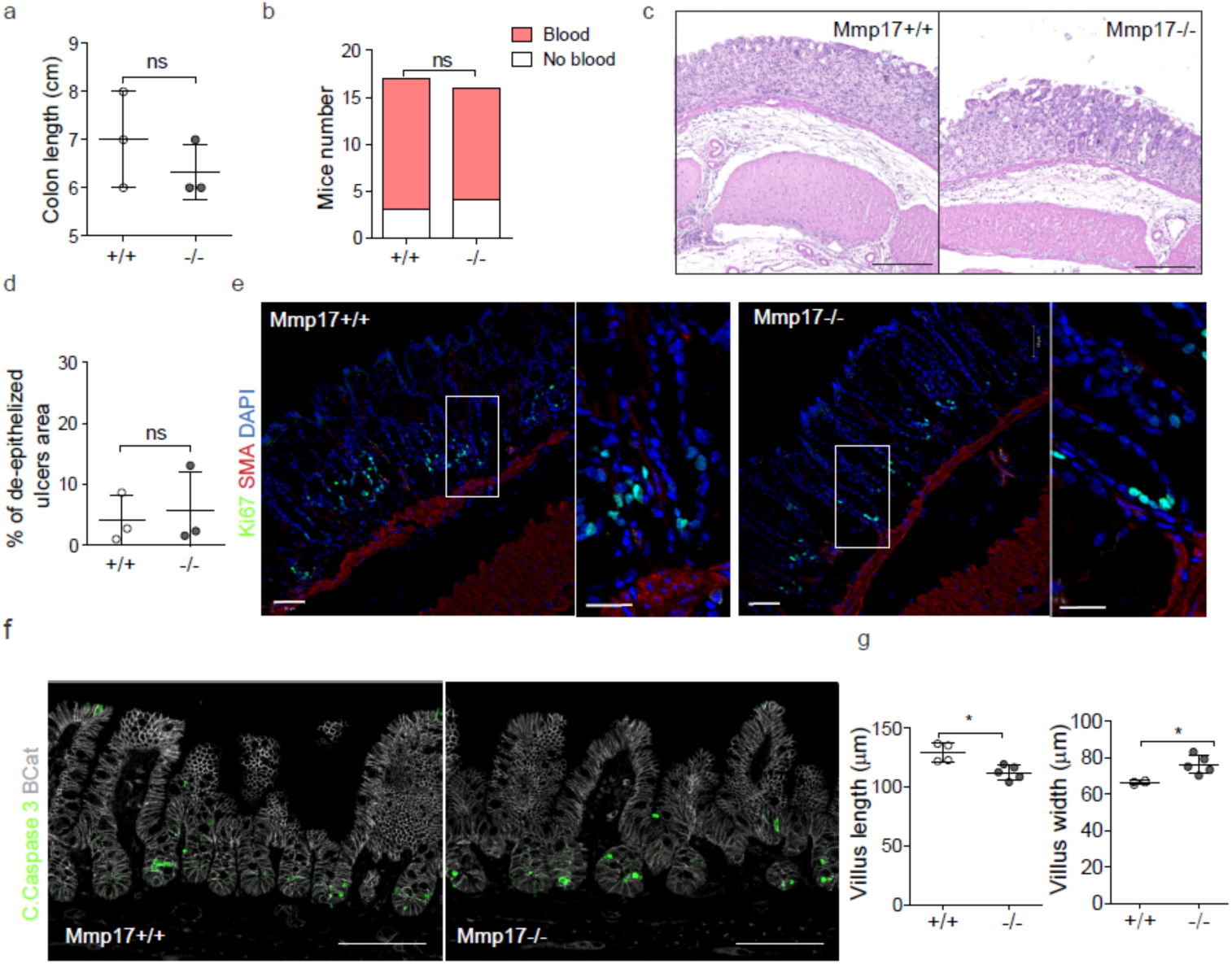
Intestinal damage models produce same injury level in WT and KO mice. **a-d.** Graphs represent colon length and presence of blood in stool at D5. n= 3 mice per genotype in a and d and 17 and 16 mice in b. c. Representative H&E picture of a transverse colon cut from mice treated with DSS 3.5% for 5 days. Scale 200 µm. n= 3 mice per genotype. d. Graph represents the presence of ulcers in the mucosa as a percentaje of the total swiss roll length at D5. n= 3 mice per genotype. e. Representative confocal images stained for Ki67 (green), SMA (red) and nuclei (blue), showing the equivalent reduction in proliferative cells at D5 of DSS treatment in WT and KO. Scale 50 µm; 25 µm in magnified view to the right. n= 3 mice per genotype. f. Representative confocal image showing Cleaved Caspase 3 staining (green) predominantly at the bottom of the crypts in ileum (24h after irradiation). βCat staining was performed to highlight epithelial cells. Scale bar 100 µm. n= 4 mice per genotype. g. Graph represents villi length and width in small intestinal tissue 3 days after irradiation. n= 10 mice per genotype in two independent experiments, analyzed 30 to 40 crypts/villi per mouse. Numerical data in a, d and g are means ± SD. Data were analyzed by Mann-Whitney test (a, d and g) and one-tailed Fisher exact test in b.

**Figure S6.**
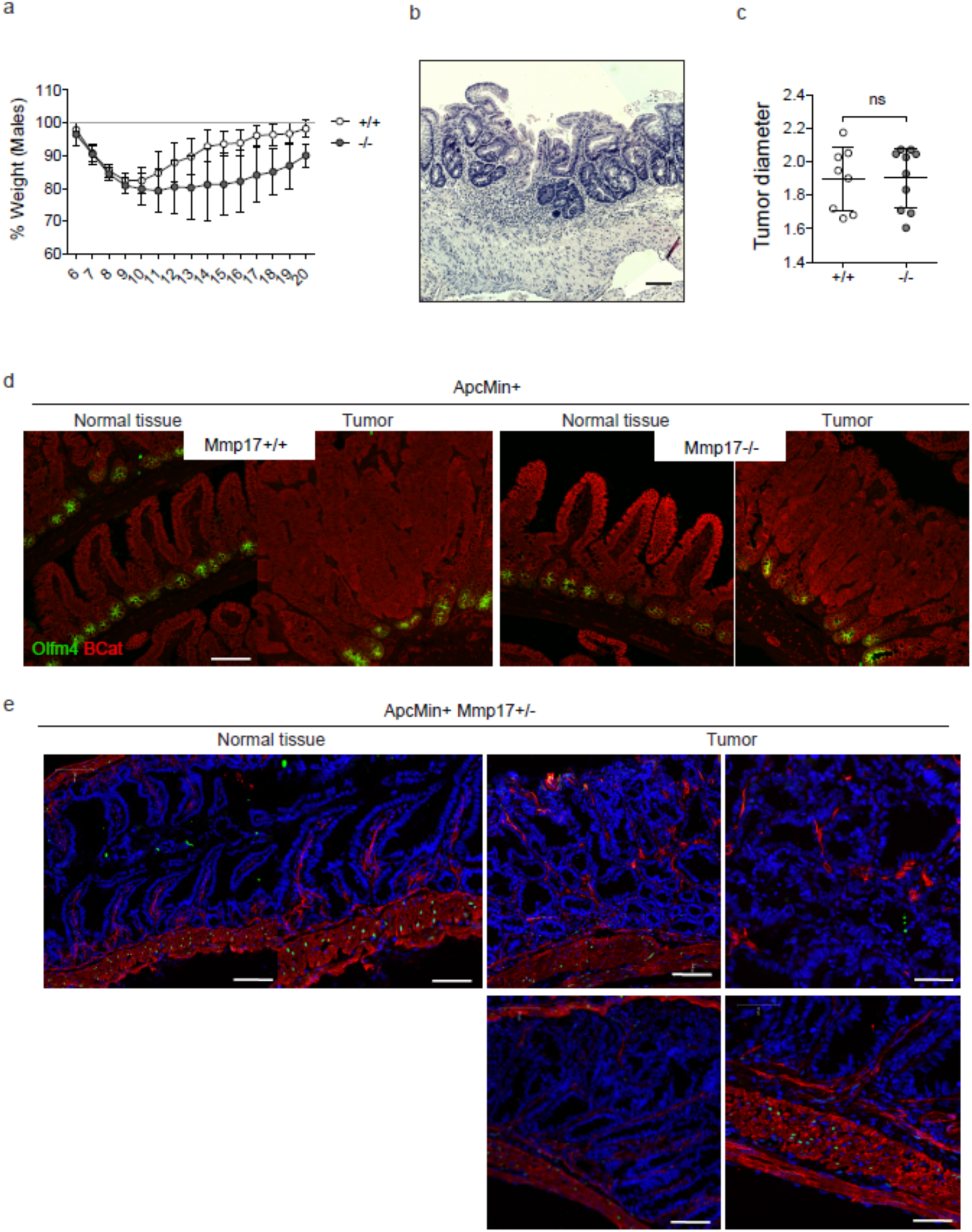
Mmp17 loss in ApcMin background predispose to tumors with no structural differences between WT and KO. **a**. Graph shows weight evolution in males after 5 days of DSS treatment. n= 4 mice per genotype. b. Representative H&E image of crypt reactive atypia in a KO mouse treated with DSS (long-term experiment). Scale bar 100 µm. n= 6-8 mice per genotype. c. Graph represents average tumor diameter (mm) in small intestines of *ApcMin+ Mmp17* WT or KO. d. Confocal maximum projection images of normal vs tumor area stained for SC marker Olfm4 (Green) and βCat (Red). Scale bar 100 µm. n= 3-5 mice per genotype. e. Representative confocal maximum projection images showing β-Gal staining (Green) in normal tissue vs tumor areas. Β-Gal signal was only found in muscle cells (SMA+, Red). Scale bar 100 µm. n= 3 mice. Data are means ± SD, numerical data in a was analyzed by two way ANOVA followed by Bonferroni post-test and t-test was applied in c.

### MATERIAL AND METHODS

#### Mice

*Mmp17-/-* mice in the C57BL/6 background have been described previously (Rikimaru et al. 2007). Mice were handled under pathogen-free conditions in accordance with CoMed NTNU institutional guidelines. Experiments were performed following Norwegian legislation on animal protection and were approved by the local governmental animal care committee. Particularly, experimental designs for DSS, irradiation procedures and ApcMin mice colony handling and tumor evaluation, were approved in advance by Norwegian authorities as stated in FOTS protocols (11842, 15888 and 17072). Mice included in these protocols were carefully monitored daily or weekly to avoid situations of moderate to high pain and comply with ethical procedures established prior to experiment development in agreement with CoMed facility at NTNU. End point protocols were applied when needed. All mice were genotyped by PCR of earclip samples using the following primers: Mt4-mmp SK1 5’-TCAGACACAGCCAGATCAGG-3’ SK2 5’-AGCAACACGGCATCCACTAC-3’ and SK3 5’-AATATGCGAAGTGGACCTGG-3’ and ApcMin: 1. 5’-TTCCACTTTGGCATAAGGC-3’ 2. 5’-GCCATCCCTTCACGTTAG-3’ 3. 5’-TTCTGAGAAAGACAGAAGTTA-3’. Experiments were conducted on mice from 8 weeks to 20 weeks of age.

#### Small intestine and colon crypt isolation

Small intestinal crypts were isolated and cultured following previously published protocol (Sato and Clevers 2013). Colon organoids were cultured according to a published method (Sato et al. 2011). Briefly, 10 first cm of the duodenum or the entire colon, were excised, flushed with cold PBS and opened longitudinally. The internal surface of the duodenum was scrapped carefully with a coverslip to remove most of the mucus and part of the intestinal villus. Small pieces of small intestine or colon (2–4 mm in length) were cut with scissors and further washed with ice-cold PBS until the supernatant was clear. Next, tissue fragments were incubated in cold 2 mmol/L EDTA chelation buffer, for 30 min (small intestine) to 1 h (colon) at 4 °C. After removal of the EDTA buffer, tissue fragments were vigorously resuspended in cold PBS (small intestine) or cold chelation buffer (colon) using a 10-mL pipette-BSA coated to isolate intestinal crypts. The tissue fragments were allowed to settle down under normal gravity for 1 minute, and the supernatant was removed for inspection by inverted microscopy. The resuspension/sedimentation procedure was repeated typically 8 times and the supernatants containing crypts (from wash 1 to 8) were collected in 50-mL Falcon tubes coated with BSA, through a 70 µm cells strainer to remove villi in case of the SI. Isolated crypts were pelleted, washed in PBS, and centrifuged at 200*g* for 3 minutes at 4°C to separate crypts from single cells. Crypts were resuspended in 10 ml basal crypt medium (BCM, advanced Dulbecco’s modified Eagle medium -F12 supplemented with penicillin/streptomycin, 10 mM HEPES, 2 mM Glutamax) to quantify its number. After centrifugation and supernatant removal, crypts were resuspended in matrigel (Corning, 734-1101) and plated in P24 well plates (150 to 250 crypts/well) or 8-well ibidi imaging µsildes (80821, Ibidi). Crypts were cultured in ENR (BCM + factors, explained below).

#### Intestinal organoid culture

Organoids were cultured in ENR medium consisting of BCM (advanced Dulbecco’s modified Eagle medium -F12 supplemented with penicillin/streptomycin, 10 mM HEPES, 2 mM Glutamax) + 1x N2 [ThermoFisher Scientific 100X, 17502048], 1x B-27 [ThermoFisher Scientific 50X, 17504044], and 1x *N*-acetyl-L-cysteine [Sigma, A7250]) and overlaid with ENR factors containing 50 ng/ml of murine EGF [Thermo Fisher Scientific, PMG8041], 20% R-Spondin-CM (conditioned medium, a kind gift from Calvin Kuo, Stanford University School of Medicine, Stanford, CA, USA), 10% Noggin-CM and, in the case of the colon, with 65% Wnt-CM (kind gifts from Hans Clevers, Hubrecht Institute, Utrecht, The Netherlands). We also made use of L-WRN (ATCC 3276) cell line conditioned media. Medium was renewed every other day. For passaging, organoids cultures were washed, and matrigel and organoids were disrupted mechanically by strong pipetting, centrifuged at 200*g*, 5 min at 4°C and resuspended in matrigel to plate in P24 wells or 8-well ibidi chambers (80821, Ibidi). In different experiments organoids were exposed to ENR, ER, EN mediums, muscle-SN, obtained as explained below, or human recombinant Periostin (50ng to 500ng) (3548-F2-050, R&D) usually for 4-5 days.

#### Intestinal muscle isolation and supernatant collection

Colon tissue was used for muscle isolation, RNA seq, WB or mass spectrometry (MS). After flushing with cold PBS, colons were open in longitudinal and dissected under a bench scope. Using a coverslip, mucosal tissue was completely removed carefully under the dissection scope and muscle was deep frozen in N2 (for RNA seq or WB analysis) or cut in 2-4 cm pieces, wash with sterile PBS twice, spin down and collected in P24 well plates. Pieces of muscle were inspected under the microscope for residual crypts or fat and further cleaned. Pieces were then incubated in DMEM-F12 for 24h before collecting the supernatant (muscle-SN). Muscle-SN was filtered (0,20 µm) and used directly + ENR factors in organoids cultures (right after splitting), sent for MS evaluation or frozen at −80°C. Organoids were exposed to muscle-SN for 4-5 days. Muscle pieces were fixed in paraformaldehyde (PFA) 4% and processed in paraffin for H&E. Tissues were inspected to confirm the absence of mucosa.

#### Quantification and imaging of organoids

P24 well plates were imaged using automatized Z-stack in EVOS2 microscope with CO2, temperature and humidity-controlled incubation chamber (Thermo-Fisher Scientific). Organoid area and classification were evaluated using a custom analysis program written in python based on opencv2 (Lindholm et al., 2020). Images where autoscaled, and a canny edge detection algorithm was run on each individual z-plane using the cv2.canny function. Small pixel groups where removed and a minimal projection of the edges was generated. The contour of objects was defined based on this image. A watershed algorithm was used to split somewhat overlapping objects from each other and the object center was defined as the pixel furthest from the edge of the object. Each object was extracted in a 120×120 image on a white background and classified as either “Junk”, “Budding” or “Spheroid” in a neural network implemented and trained using Tensorflow and Keras. A custom visual classification editor was then used to correct locations of organoid centers and the classification group. The order of images was randomized, and treatment information hidden during the classification correction. The images then went through a second segmentation step where the segmentation was re-done as described above. The objects of contours with multiple centers were then split apart with a watershed algorithm that used the corrected organoid centers to split overlapping organoids. Organoids formation efficiency was calculated by manually counting the number of successful organoids 24, 48 and 72h after crypt isolation in EVOS2 bright field pictures. Organoids size was evaluated from day 1 to 5. For muscle-SN experiments quantification was performed at day 4 and at day 3 in POSTN treatments.

#### Immunofluorescence staining in organoids and imaging

For immunofluorescent labeling and imaging, organoids were grown in 70% Matrigel-30% ENR on eight-chamber Ibidi μslides (80821, Ibidi). Organoids were fixed in PBS containing 4% paraformaldehyde (pH 7.4) and 2% sucrose for 30-45 min, permeabilized, and blocked in PBS-Triton X-100 0.2%, 2% normal goat serum (NGS), 1% BSA Glycine 100 µM for 1h at RT. Next, organoids were incubated with primary antibodies against the following antigens diluted in PBS-TX100 0.2% + BSA 0.5% + NGS 1%: Ki67 (1:200, rabbit monoclonal antibody (mAb), Invitrogen, MA5-14520), β-catenin (1:200, mouse mAb, BD Biosciences, 610154), and YAP (1:100, rabbit mAb, Cell Signalling, 14074) overnight at 4°C with slow agitation. Organoids were washed in PBS containing 0.1% Tween20 and incubated overnight in the same buffer at 4°C with the appropriate Alexa Fluor secondary antibody (1:500) along with Hoechst 33342 (1:10,000). Organoids were washed with PBS buffer with 0.1% Tween 20 and mounted using Fluoromount G (ThermoFisher Scientific, 00-4958-02). Organoids were imaged in a Zeiss Airyscan confocal microscope, using a 10x and 20x objective lens. Images were analyzed using Zen black edition software (Zeiss) and maximal projections are shown.

#### RNA seq of organoids and tissue

Pieces of clean colon muscle, colon crypts (obtained using colon crypt isolation protocol previously described), or pooled organoidswere used for RNA isolation. Tissues were first placed in lysis buffer (RLT, provided in RNeasy® Mini Quiagen Kit, 74104) and disrupted with sterile ceramic beads (Magna Lyser green beads tubes 03358941001) using a Tissue Lyser (FastPrep-24™, SKU 116004500), with two rounds of 6500 rpm for 30 seconds each, with care taken to maintain the sample cold. Organoid wells were first disrupted by strong pipetting and pelleted before lysis. RNA isolation was performed following manufacturer instructions (RNeasy® Mini Kit, Qiagen, 74104, for tissue and Direct-zol™ RNA MiniPrep, BioSite R-2052, for organoids). RNA was quantified by spectrophotometry (ND1000 Spectrophotometer, NanoDrop, Thermo Scientific) and 25 µl at a 50 ng/µl concentration of RNA were used for RNA seq. Library preparation and sequencing for tissue RNA seq was performed by NTNU Genomic Core facility. Lexogen SENSE mRNA library preparation kit was used to generate the library and samples were sequenced at 2×75 bp paired end using Illumina NS500 flow cells. Library preparation and sequencing for organoid RNA seq was performed by Novogene (UK) Co. NEB Next® Ultra™ RNA Library Prep Kit was used to generate the library and samples where sequenced at 150×2 bp paired end using a Novaseq 6000 (Illumina). The STAR aligner was used to align reads to the Mus musculus genome build mm10(Dobin et al. 2013), (Frankish et al. 2019). featureCounts was used to count the number of reads that uniquely aligned to the exon region of each gene in GENCODE annotation M18 of the mouse genome(Liao, Smyth, and Shi 2014). Genes that had a total count of less than 10 were filtered out. DESeq2 with default settings was used to do a differential expression analysis(Love, Huber, and Anders 2014). Heatmaps were generated using the R-package pheatmap(Kolde 2015). Principal component analysis (PCA) analysis was done with the scikit-learn package using the function sklearn.decomposition.PCA(Pedregosa et al. 2011). Gene set enrichment analysis (GSEA) was performed on the full list of genes from differential expression sorted by log2 fold change and with log2 fold change as weights. GSEA was run with the R package clusterProfiler using 10000 permutations and otherwise default settings(Yu et al. 2012). A list of the top 250 genes upregulated in crypts was used for Enrichr analysis(Chen et al. 2013; Kuleshov et al. 2016).

All raw sequencing data is available online through ArrayExpress: WT and KO smooth muscle and crypt RNA seq: E-MTAB-9180; ENR vs MuscleSN treated organoids RNA seq: E-MTAB9181.

#### Mass spectrometry of muscle-SN

Muscle supernatants were collected as stated above. Proteins were reduced with 4 mM DTT at room temperature for 1 hour, and alkylated with 8 mM iodoacetamide at room temperature for 30 min in the dark, after which additional 4 mM DTT was added. A first digestion was carried out with 40 ng Lys-C at 37 °C for 4 h. The samples were diluted four times and further digested with 40 ng trypsin at 37 °C overnight. Digested protein were desalted using Sep-Pak C18 cartridges (Waters), dried by vacuumcentrifuge and stored at−20 °C for further use.

Before analysis the peptides were desolved in 2% formic acid. The digested samples were injected on an Orbitrap Q Exactive HF spectrometer (Thermo Scientific) connected to a UHPLC 1290 system (Agilent). Reconstituted peptides were trapped on a double-fritted (Dr Maisch Reprosil C18, 3µM, 2cm x 100µM) precolumn for 5min in solvent A (0.1% formic acid in water) before being separated on an analytical column (Agilent Poroshell, EC-C18, 2.7µM, 50cm x 75µM). Solvent A consisted of 0.1% formic acid, solvent B of 0.1 % formic acid in 80% acetonitrile. A 95 min gradient of 13–44 % buffer B followed by 44–100% B in 3 min, 100% B for 1 min was applied. MS data were obtained in data-dependent acquisition mode. Full scan MS spectra from m/z 375 – 1600 were acquired at a resolution of 60.000 to a target value of 3×106 or a maximum injection time of 20ms. The top 15 most intense precursors with a charge state of 2+ to 5+ were chosen for fragmentation. HCD fragmentation was performed at 27 normalized collision energy on selected precursors with 16s dynamic exclusion at a 1.4m/z isolation window after accumulation to 1×105 ions or a maximum injection time of 50ms. Tandem mass spectrometry (MS/MS) spectra were acquired at a resolution of 30.000.

Proteins IDs and intensity values (abundance) are represented in Table 1.

#### Tissue immunostaining and immunohistochemistry

Intestines were open in longitudinal, flushed and rolled into “Swiss rolls” as previously described(Moolenbeek and Ruitenberg 1981). Tissues were then fixed in PFA 4% 16h and, in the case of frozen samples, included in sucrose 2% from 3h to over night. Swiss rolls were then included in paraffin or in OCT compound for frozen sectioning. Transversal cuts of swiss rolls were used in all experiments. For H&E staining, paraffin sections of 4 µm were subjected to normal deparaffination and hydration. For LacZ staining, frozen sections were stained following β-gal staining kit indications (K1465-01, Fisher Scientific). Tissue sections were imaged using brightfield microscope EVOS2 with 10 or 20X objective lens and tile scan for the visualization of the complete swiss roll was performed when needed. For immunofluorescence, sections of 4 to 7 µm were incubated in blocking buffer for 1h (PBS-Tx100 0.3%, NGS 5%, BSA 2%, Glycine 100 µM), followed by primary antibody incubation overnight. The following antibodies were used for immunofluorescence: anti-βGalactosidase (βGal, Frozen sections, Dil 1:100, Rabbit polyclonal antibody, Abcam, ab4761), anti-Ki67 (Paraffin, 1:200, rabbit monoclonal antibody, Invitrogen, MA5-14520) anti-SMAD4 (Frozen sections, methanol 10min −20C, Dil 1:400, Cell signaling, 46535), anti-βcatenin (Paraffin, 1:200, Mouse mAb, BD Biosciences, 610154), anti-Olfm4 (Paraffin, 1:200, Rabbit mAb, Cell signaling, 39141), anti-cleaved caspase 3 (Paraffin, Dil 1:200, Rb pAb, Cell signaling, 9661) and anti-pSMAD1/5/9 (Frozen sections, 10 µm, Dil 1:800, 13820, Cell Signaling, following TSA amplification, NEL744001KT, Perkin Elmer. following previous specifications(Huycke et al. 2019)). After washing the slides in PBS-Tween 20 2%, tissues were incubated with the appropriate Alexa Fluor secondary antibody, Anti-SMA-Cy3 directly labeled antibody (1:500, mouse mAb, Sigma-Aldrich C6198) and Hoechst 33342 (1:10,000). Tissue sections were imaged with a Zeiss Airyscan confocal microscope, using a 10x and 20x objective lens. Images were analyzed using Zen black edition software (Zeiss) and maximal projections are shown except for SMAD4 staining (single plane). Tile scans of swiss rolls were performed when required. Percentage of βGal positive cells in the muscle was calculated manually in Zen software by counting the total number of nuclei in muscularis mucosa or circular/longitudinal muscle vs βGal positive nuclei in these tissues.

#### Dextran Sulfate Sodium (DSS) colon injury

Experimental epithelial colon injury was induced in 8 to 10 weeks mice by supplying 3.5% dextran sodium sulfate (DSS) (MP Biomedicals, 0216011010) in drinking water ad libitum during 5 days. Before (Day 0) and during experimental development (up to day 7 in short protocol or up to 19 days in long-term protocol), mice were monitored daily for signs of stress, pain, body weight loss and presence of blood in stool (detected by hemoFEC, Cobas, 10243744). At day 5, DSS was replaced by regular water to allow epithelial renewal. Control mice to evaluate DSS damage level were euthanized at Day5. For the long-term protocol, weight recovery determined end point (14 days for females and 19 days for males). Colon tissue was then harvested, measured, imaged, fluxed with cold PBS, open in longitudinal and processed for paraffin or OCT, as indicated above. Paraffin sections were stained for H&E, or Ki67 and SMA. H&E sections were imaged on EVOS2 (tile scan of complete swiss rolls) and genotype-blind analyzed for signs of injury. Crypt loss, presence of ulcered tissue in the mucosa, signs of immune infiltrate in mucosa and edema were evaluated to create an injury level profile for each sample (Injury Score). The % of de-epithelialized mucosa was calculated using Fiji (Image J) as the total length of intestinal surface devoid of epithelium vs total length of swiss roll. The % of Ki67+ cells was obtained by quantifying Ki67 positive nuclei in mucosa vs total nuclei in mucosa using Cell Profiler. Long-term DSS samples were evaluated by a pathologist to determine presence of reactive atypia.

#### Irradiation

8 week-old male mice were subjected to whole body 10Gy irradiation under anesthesia. Weight loss was evaluated at Day 0 and during the length of the experiment (3 days). Animals were carefully checked daily for signs of pain or stress. At end point, mice were euthanized and small intestine and colon were harvested, flushed with cold PBS, and processed in swiss rolls for paraffin. H&E sections were imaged and evaluated blindly for signs of impaired recovery of the mucosal tissue. Parameters as crypt loss, immune infiltrate, crypt length and villus length and width were evaluated to rate the injury score. Villus length and width were measured using Fiji (Image J). We evaluated caspase 3 levels through staining at Day 1 as a positive control for irradiation.

#### Mmp17-ApcMin tumor evaluation

The presence and number of intestinal tumors were evaluated in 18 to 20 weeks old *ApcMin+ Mmp17+/+*, *Mmp17-/-* or *Mmp17+/-* (females and males) mice. Mice were weekly monitored for weight loss and signs of pain, to avoid situations of moderate to high pain. At end point, mice were euthanized and small intestinal tumors were visualized and counted. Brightfield pictures were taken to double check tumor number and quantify tumor size (Fiji, Image J). Tissues were fixed as swiss rolls overnight and processed in paraffin or for OCT as described above 5 um microtome cuts were stained for H&E and/or for immunostaining. Immunostaining was performed as previously described in paraffin for βCatenin and Olfm4 staining. For the evaluation of *Mmp17* expression in tumors, *Mmp17+/-* tissues were processed in OCT and stained for βGal as stated above.

#### Protein extraction and western blot analysis

Intestinal tissue (muscle or mucosa tissue isolated as previously described) was disrupted using a tissue lyzer and ceramic beads (Magna Lyser green beads tubes 03358941001) in cold regular RIPA buffer (+ protease and phosphatase inhibitors). Proteins (20 to 100 µg) were resolved by 7-12% SDS-PAGE in reducing conditions and transferred onto nitrocellulose membranes using iBlot2^TM^ dry blotting technology (Thermo Fisher Scientific). Antibodies against SMAD4 (rabbit mAb, Cell signaling, 46535), POSTN (mouse mAb, SAB4200197, MERK, Sigma-Aldrich), β-tubulin (ab6160, Abcam,) and GAPDH (mouse mAb, Abcam, ab125247) were used at 1:1000 dilution (4°C overnight). Immunoreactive proteins were visualized with corresponding fluorochrome-conjugated secondary antibodies (680 or 800 ODYSSEY IRDye®) and recorded by Licor Odyssey technology. Quantification of western blot bands was performed using ODYSSEY software (LI-COR Biosciences) and normalized to GAPDH or β-tubulin levels.

#### In vitro digestion and western blot

Human recombinant POSTN (250 ng to 1 µg; hrPOSTN, 3548-F2-050, R&D) and the catalytic domain of human recombinant MMP17 (250 ng) (hrMMP17, P4928, Abnova) were incubated in digestion buffer (50 mM Tris-HCl, 10 mM CaCl2, 80 mM NaCl, [pH7.4]) for 2 hours at 37°C. Samples were separated by 12% SDS-PAGE and transferred to nitrocellulose membranes. Full length hrPOSTN and fragments were detected with an anti-POSTN monoclonal mouse antibody (SAB4200197, MERK, Sigma-Aldrich). An anti-mouse secondary antibody was used to visualize the different fragments.

#### In situ hybridization (ISH) of intestinal tissues

ISH using RNAscope technology (ACD, Bio-Techne) was performed using freshly cut paraffin sections of intestinal swiss rolls of *Mmp17+/+* and *Mmp17-/-* mice. Chromogenic RNAscope of *Lgr5* (Mm-Lgr5, 312171) and *Olfm4* (Mm-Olfm4, 311831) in colon and small intestine was performed following manufacturer indications (RNAscope® 2.5 HD Assay-BROWN). Sections were imaged using bright field automatized microscope EVOS2 (tile scan). Quantification of positive Lgr5 cells was performed using Fiji (Image J); total area of positive Lgr5 crypts was normalized to tissue length. Immunofluorescence RNAscope (FISH) was performed following manufacturer protocol (RNAscope® Multiplex Fluorescent v2) and the following probes were used: *Mmp17* (Mm-Mmp17-C4 ACD design of NM_011846.5), *Grem1* (Mm-Grem1-C3, 314741), *Grem2* (473981-C2), *Chrdl1* (Mm-Chrdl1, 442811). Zeiss Airyscan scope images are maximal projections of the different channels taken with 10X and 20X objectives.

#### Statistical analysis

The statistical analysis performed in each case is explained in detail in the corresponding figure legend, together with the n and the times the experiment was performed. Data were analyzed by two-tailed Student’s t-test or by one- or two-way ANOVA followed by Tukey multiple comparison test for data with Gaussian normal distribution. Non-normal distributions were analyzed by Mann-Whitney test. The one-tailed Fisher’s exact test was used to analyze the incidence of blood in stool or the presence of reactive atypia. Statistical tests were conducted with Prism 5 software (GraphPad Software, Inc.). Data are presented as mean ± S.D. and differences were considered statistically significant at p < 0.05: *p value < 0.05, ** p value < 0.01, *** p value < 0.001 and **** p value < 0.0001.

